# Quantifying and visualising divergence between pairs of phylogenetic trees: implications for phylogenetic reconstruction

**DOI:** 10.1101/227942

**Authors:** Martin R. Smith

## Abstract

Tree topologies are a primary output of phylogenetic analysis. It is important to understand how such outputs are influenced by the choice of phylogenetic method, and the nature and quality of input data. This often entails the measurement of how closely an output corresponds to an ideal tree topology. The symmetric difference (partition) metric is the only widely-used measure that is defined for trees that contain polytomies, but expresses undesirable behaviour in certain situations.

Here I propose a divergence metric for unweighted tree topologies based on quartet statements, which measures the topological information common to two trees. This metric reflects the contributions of both accuracy and precision to tree quality; these components can be decomposed through the use of ternary diagrams. This approach is implemented in a new package for the R statistical environment, and applied to a recent controversy, where it provides a new perspective on the relative merits of Bayesian and parsimony approaches to morphological phylogenetics.

## INTRODUCTION

Because phylogenetic trees are not vectors, there is no natural scale against which they can be compared (Penny & Hendy, 1985). Whilst it is possible to measure distances between edge-weighted trees in multi-dimensional space (Billera, Holmes, & Vogtmann, 2001; Owen & Provan, 2011), not all analyses generate meaningful edge lengths, meaning that their output can be difficult to compare.

This is unfortunate, as tree comparison underpins many fundamental considerations in phylogenetic analysis. Tree similarity has been used to determine the effect of missing data on phylogenetic analysis (Guillerme & Cooper, 2016), the impact of fossilization on phylogenetic results (Pattinson, Thompson, Piotrowski, & Asher, 2015; Sansom & Wills, 2013), the reliability of morphological data from different anatomical regions (Clarke & Middleton, 2008; Sansom, Wills, & Williams, 2017), and the most suitable method for morphological analysis (Congreve & Lamsdell, 2016; O’Reilly et al., 2016).

All existing measures of tree similarity have limitations.

The SPR distance measures the number of subtree pruning and regrafting rearrangements necessary to transform one of a pair of trees into the other (Penny & Hendy, 1985). Its limited range of integer values do not allow it to distinguish similar trees (one tip moved one edge) from more dissimilar trees (a large clade moved a long distance), and it is only defined for pairs of bifurcating trees.

The path difference (Steel & Penny, 1993) counts differences in the number of edges that must be traversed to travel between each of the pairs of tips in a tree. The square of the difference in path length between the two trees is calculated for each pair of tips, and the square root of the sum of these quantities gives the path difference. The metric has high resolution, and behaves as expected in all situations, though the interpretation of its value is not straightforward, and it is only defined for pairs of bifurcating trees.

The symmetric difference measure (Penny & Hendy, 1985), a refinement of the Robinson-Foulds partition metric (Robinson & Foulds, 1981), is not restricted to bifurcating trees. It counts the proportion of edges (i.e. bipartitions or splits) present in each tree that are absent in the other. The measure has limited resolution, and can reach its maximum value when trees differ only in the position of a single tip (Steel & Penny, 1993). Worse, moving a single tip to a particular location can generate a higher distance metric than moving both that tip and its immediate neighbour to the same location. These counterintuitive properties mean that the metric offers a biased measure of tree similarity.

Finally, the quartet metric counts the number of four-taxon trees that are present as subtrees of each tree under consideration (Estabrook, McMorris, & Meacham, 1985; Steel & Penny, 1993). This method has a high resolution, allowing it to distinguish between relatively similar differences; and it does not exhibit the anomalous behaviour of the symmetric difference measure.

The method can be adapted in various ways to address pairs of trees that contain polytomies. If the trees each contain *n* tips, each of the *Q* = *_n_C*_4_ combinations of four taxa (i.e. quartets) will satisfy one of five conditions (Estabrook et al., 1985): the quartet may be resolved in the same way (*s*) or a different way (*d*) on each tree; resolved in tree 1 only (*r*_1_); resolved in tree 2 only (*r*_2_); or unresolved in both trees (*u*). The proportions of quartets satisfying each of these conditions can be combined in various ways in order to capture different aspects of how trees agree, disagree, or differ in resolution (Estabrook et al., 1985).

One approach to combining these variables is to emphasize accuracy, measured by the number of incorrect quartets (i.e. “type II error over type I error” in Congreve & Lamsdell, 2016). Such a measure is unsatisfactory, as no tree can score better than a single polytomy that resolves no quartets incorrectly). The other extreme is to emphasize precision (Carpenter, 1988), which represents a preference for inaccuracy over ignorance, as better-resolved trees will typically contain a higher proportion of erroneous nodes.

The compromise position recognises accuracy and precision as two different but fungible components of the information held by a tree (MacKay, 1953). We desire a measure that allocates higher values to trees that more informatively describe a comparison tree, and whose value will not be affected by an information-neutral barter between precision and accuracy, for example by collapsing or resolving weakly supported nodes.

## METHODS

If each quartet is taken to represent a single unit of information, then such a metric can be defined by calculating the number of quartet operations necessary to change from one tree into the other: each of the *d* + *r*_1_ quartets that are present in the first tree must be unpicked, and each of the *d* + *r*_2_ quartets unique to the second tree must be forged. (2*d* + *r*_1_ + *r*_2_) / *2Q* provides an information-based measure of tree dissimilarity that reflects both accuracy and precision.

This measure is mathematically equivalent to the normalised symmetric difference metric, though it counts quartets in place of partitions as the unit of information. I call the measure the quartet divergence, by analogy with the Kullback-Leibler divergence (Kullback & Leibler, 1951) – though quartets are not independent so do not satisfy the statistical properties of Shannon-Weiner information.

A high value of this metric may represent high resolution (but some misinformation) or high accuracy (tempered by low resolution). The distinction between precision and accuracy can be instructive (e.g. Brown, Parins-Fukuchi, Stull, Vargas, & Smith, 2017), but is difficult to visualise. Because precision affects a tree’s resolution and similarity score, plots that use these terms as their axes (e.g. O’Reilly et al., 2016; Puttick et al., 2017), will be dominated by autocorrelation, making them difficult to interpret. I instead employ a ternary plot with corners corresponding to the proportion of quartets (or partitions) that are the same in both trees (*s*), different in both trees (*d*), and unresolved in at least one tree (*r*_1_ + *r*_2_ + *u*).

### Implementation

The new R package `SlowQuartet` uses explicit enumeration to calculate the condition of each quartet, an O(*n*^4^) approach that is only practical for trees with up to a few dozen tips. A quicker O(*n* log *n*) solution (Sand et al., 2014) is applied to pairs of bifurcating trees. Ternary plots are produced using the Ternary package (Smith, 2017).

### Case study

I used these approaches to visualise the results of a simulation study that evaluates different methods of phylogenetic reconstruction (Congreve & Lamsdell, 2016). These authors simulated 55-character matrices from a 22-tip bifurcating tree using a Markov *k*-state 1 parameter model (Mk1, Lewis, 2001) with a gamma parameter (Wright & Hillis, 2014). I used TNT v1. 1 (Goloboff, Farris, & Nixon, 2008) to conduct parsimony search with equal and implied weights, using the parsimony ratchet (Nixon, 1999) and sectorial search (Goloboff, 1999) heuristics (search options: `xmult:hits 20 level 4 chklevel 5 rat10 drift10`). I took a strict consensus of all optimal trees obtained under equal weights, and under implied weights (Goloboff, 1993) at concavity constants of *k* = 1, 2, 3, 5 and 10. For each dataset I generated a further strict consensus of all trees that were optimal under any of the concavity constants 2, 3, 5 and 10, but not *k* = 1 – which represents the extreme philosophy (Smith & Caron, 2015) that each step beyond the first makes a negligible contribution to tree score.

I also generated majority-rule consensus trees in MrBayes 3.2.2 (Huelsenbeck & Ronquist, 2001) using an Mk model (Lewis, 2001), with rates distributed according to a gamma parameter. I conducted four runs of four chains for 10 000 000 generations, sampling every 10 000 generations and discarding the first 40% of samples as burn-in (Topology parameter: 0.999 < PSRF < 1.001; ESS > 400). Scripts are provided in the Supplementary Information.

For each analytical configuration, I generated 20 further trees by progressively lowering the resolution of the most resolved tree. Under the Mk model, I collapsed clades with a posterior probability of > *x*%, with *x* varying uniformly from 50–100. In parsimony analyses, strict consensus trees were produced including trees that were suboptimal by *x* steps (TNT command `subopt bbreak;`), with *x* corresponding to the integers 1..20 for implied weights, and drawn from a logarithmic distribution (0.73^0…19^, 2.5×10^−3^→1×10^0^) for implied weights.

To generate summary statistics, *s*, *d*, and *r*_2_ were calculated for each tree relative to the bifurcating generative tree (*r*_1_ = *u* = 0), and the mean of each of parameter was calculated for each analysis at each value of *x*.

## RESULTS

There is no significant difference (at *p* = 0.01) between the quartet divergence of the best trees generated by the *Mk* model or implied weights (*k* ∈ {2, 3, 5, 10}), but the best trees generated by equal weights, or implied weights with *k* = 1, are significantly worse than those produced by the other methods (Figure 1).

**Figure 1.**
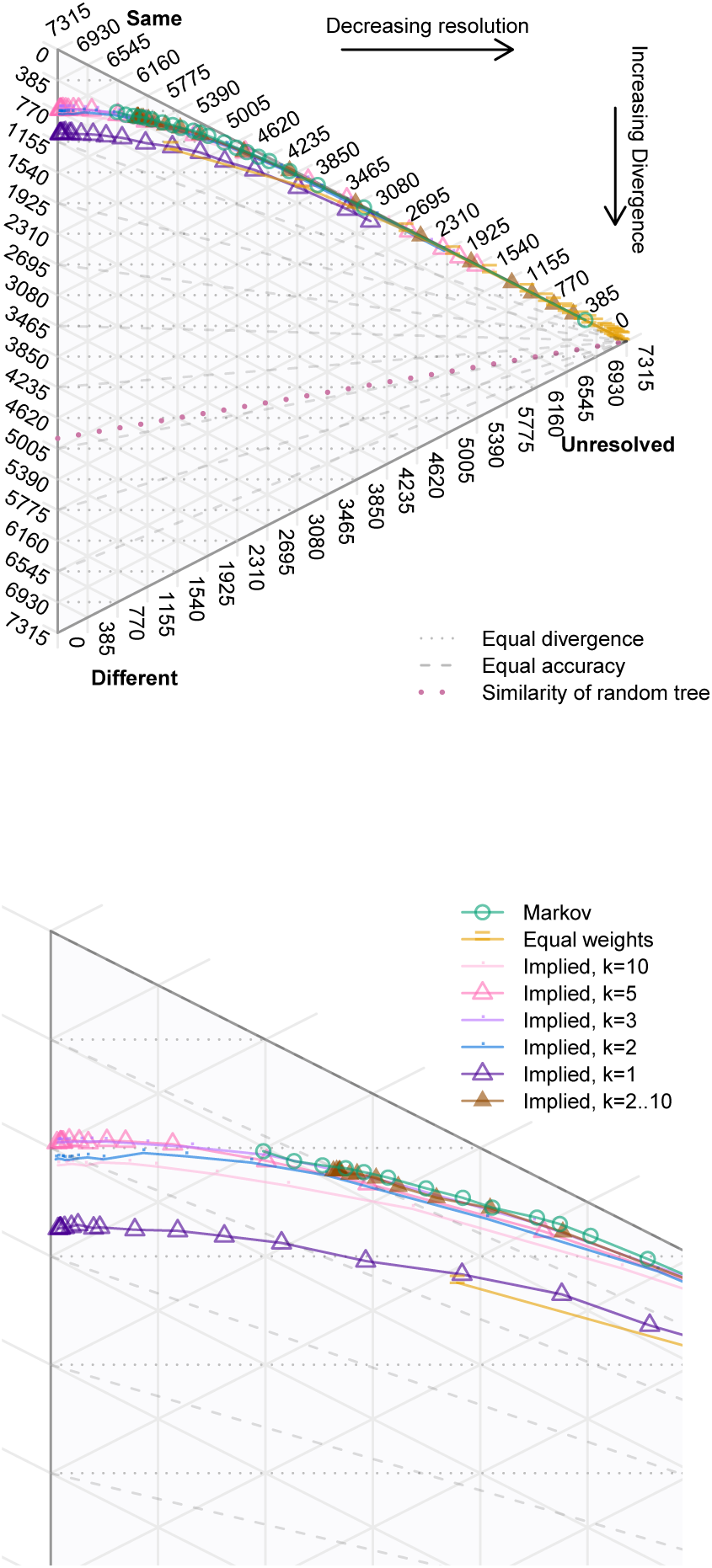
Quartet divergence, accuracy and precision of trees recovered by different analytical methods. Average number of quartets, across all 100 simulated datasets, that are unresolved, the same as the generative tree, or resolved differently to the generative tree. Each marked point corresponds to the strict consensus of all trees that are suboptimal by a given amount under the specified given method.

Implied weights generated the highest precision, but not the highest accuracy (consistent with O’Reilly et al., 2016). Collapsing the least-supported nodes initially increases the accuracy (i.e. proportion of correctly to incorrectly resolved nodes), leading to a trivial increase in the overall informativeness of the tree. After a point, however, the gain in accuracy no longer offsets the information lost by collapsing nodes, and the tree diverges increasingly from the generative tree. The lower resolution of the equal weights and Bayesian results means that they do not experience this initial increase in tree quality: collapsing nodes immediately increases divergence from the generative tree.

Similar results occur if partitions are used instead of quartets to calculate tree divergence (Figure 2). The results of implied weights (*k* ∈ {2, 3, 5, 10}) and Bayesian analysis are statistically indistinguishable (at *p* = 0.01), whilst equal weights and implied weights with *k* = 1 generate trees that are significantly worse. Relative to the quartet measure, however, collapsing nodes more readily improves the partition score of a tree; optimal trees are obtained once relatively many nodes have been collapsed – indicating that the partition measure emphasizes accuracy over precision. These overall patterns and relationships continue to hold if datasets with a low consistency index are excluded.

**Figure 2.**
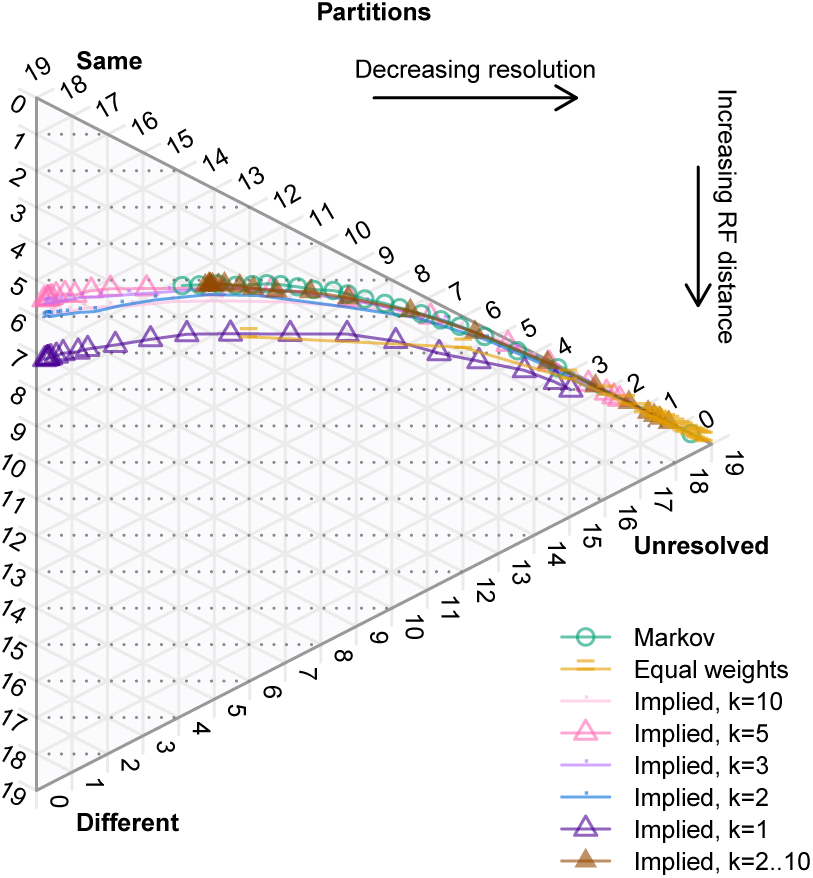
Partition resolution of trees recovered by different analytical methods. Average number of quartets, across all 100 simulated datasets, that are unresolved, the same as the generative tree, or resolved differently to the generative tree. Each marked point corresponds to the strict consensus of all trees that are suboptimal by a given amount under the specified given method.

## DISCUSSION

The strict preference for accuracy (i.e. number of incorrect nodes) over resolution espoused by Congreve and Lamsdell (2016) can be visualised as a preference for trees that are closest to the upper right edge of the ternary space (Figure 3). This philosophy illustrates the danger of examining only the most resolved tree that a given method can produce (as discussed by Brown et al., 2017): even if the best-resolved tree of equal weights falls closer to this edge than the best-resolved trees under equal weights, collapsing nodes to polytomies causes all methods to approach the optimal score of zero incorrect nodes.

**Figure 3.**
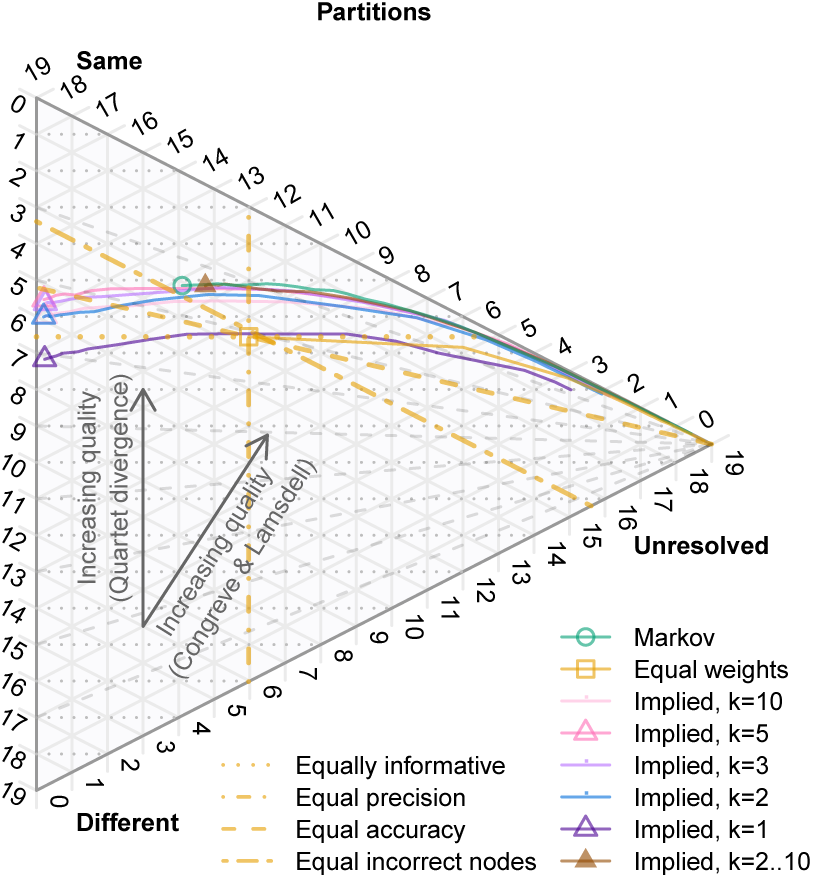
Four measures of tree quality. Congreve and Lamsdell (2016) equate tree quality with the number of incorrect nodes. Looking only at most resolved trees, equal weights outperforms implied weights on this measure. But then under any measure of accuracy that neglects precision, trees can be improved by continuing to collapse their least-supported nodes, so the optimal tree is entirely unresolved. Favouring precision alone does not distinguish between methods that can produce perfectly-resolved trees, regardless of whether such trees are accurate. The most informative (vertically highest) trees, in contrast, strike a balance between precision and accuracy.

Divergence metrics offer a more meaningful comparison of methods, and demonstrates that at any given level of precision, equal weights and the extreme implementation of implied weights with *k* = 1 are significantly less accurate than other methods.

In general, the increase in accuracy attainable by intelligently reducing resolution (Goloboff, 1995; Salisbury, 1999) more than compensates for any reduction in precision, up to a point: though the partition metric differs from the quartet divergence metric in where this optimal trade-off lies. As a rule of thumb, collapsing weakly-supported nodes until trees reach an equal resolution to Bayesian results allows implied weights parsimony to generate trees that have an equivalent accuracy to Mk trees, removing a reason to prefer Bayesian analysis to parsimony (cf. O’Reilly et al., 2016). Even more accuracy might be gained by collapsing nodes if other measures of node support (e.g. Giribet, 2003; Goloboff et al., 2003) are employed.

In conclusion, the quartet divergence metric produces notably different results to the partition metric of tree distance, with different implications for best practice. Ternary plots provide a means to visualise the contributions of accuracy and precision to tree quality, illustrating that perceived inaccuracy of implied weighting trees is simply a function of the greater precision that this method can, but need not, attain

## ACKNOWLEDGEMENT

The TNT software is supported by the Willi Hennig Society.

## DATA AVAILABLITY

The `SlowQuartet` package is available at https://github.com/ms609/SlowQuartet. Its vignettes depict analytical results for each individual Congreve & Lamsdell dataset. Supplementary data files have been uploaded to FigShare, 10.6084/m9.figshare.5659195.

